# Dissociating Contributions of Theta and Alpha Oscillations from Aperiodic Neural Activity in Human Visual Working Memory

**DOI:** 10.1101/2024.12.16.628786

**Authors:** Quirine van Engen, Geeling Chau, Aaron Smith, Kirsten Adam, Thomas Donoghue, Bradley Voytek

## Abstract

While visual working memory (WM) is strongly associated with reductions in occipitoparietal alpha (8-12 Hz) power, the role of frontal midline theta (4-7 Hz) power is less clear, with both increases and decreases widely reported. Here, we test the hypothesis that this theta paradox can be explained by non-oscillatory, aperiodic neural activity dynamics. Because traditional time-frequency analyses of electroencephalography (EEG) data conflate oscillations and aperiodic activity, event-related changes in aperiodic activity can manifest as task-related changes in apparent oscillations, even when none are present. Reanalyzing EEG data from two visual WM experiments (n = 74, of either sex), and leveraging spectral parameterization, we found systematic changes in aperiodic activity with WM load, and we replicated classic alpha, but not theta, oscillatory effects after controlling for aperiodic changes. Aperiodic activity decreased during WM retention, and further flattened over the occipitoparietal cortex with an increase in WM load. After controlling for these dynamics, aperiodic-adjusted alpha power decreased with increasing WM load. In contrast, aperiodic-adjusted theta power appeared to increase during WM retention, but because aperiodic activity reduces more, it falsely appears as though theta “oscillatory” power (e.g., total band power) is reduced. Furthermore, only a minority of participants (31/74) had a detectable degree of theta oscillations. These results offer a potential resolution to the theta paradox where studies show contrasting power changes. Additionally, we have identified novel aperiodic dynamics during human visual WM.

**Significance statement:** Working Memory (WM) is our ability to hold information in mind without it being present in our external environment. Years of research focused on oscillatory brain dynamics to discover the mechanisms of WM. Here, we specifically look at oscillatory and non-oscillatory, aperiodic activity as measured with scalp EEG to test their significance in supporting WM. We challenge earlier findings regarding theta oscillations with our analysis approach, while replicating alpha oscillation findings. Furthermore, aperiodic activity is found to be involved in WM, over frontal regions in a task-general manner, and over anterior regions this activity is reduced with an increase in the number of remembered items. Thus, we have identified novel aperiodic dynamics during human visual WM.

## 1. Introduction

We commonly encounter situations where we need to remember the identity of objects in our environment, ranging from simple tasks like cooking, to the games we play, to more complex interactions. Our visual working memory (WM) allows us to actively maintain information in mind for a brief period of time; it guides our behavior to execute these tasks (Baddeley, 1992; D’Esposito & Postle, 2015) and is characterized by its limited capacity (Cowan, 2010; Miller, 1956; Oberauer et al., 2016). Using electroencephalography (EEG), systematic changes in neural activity are observed while participants maintain items in memory, even in the absence of continuing sensory input.

Two oscillatory bands likely play an important role in WM: frontal midline theta (4-7Hz, hereafter “theta”) (Berger & Sauseng, 2022; Hsieh & Ranganath, 2014; Jensen & Tesche, 2002), and occipitoparietal alpha (8-12Hz, hereafter, “alpha”) (D’Esposito & Postle, 2015; Gevins & Smith, 2000; Halgren et al., 2002; Pavlov & Kotchoubey, 2022). Alpha power decreases are observed with increasing WM load (Fukuda et al., 2015, 2016), whereas hemispheric load differences have inconsistent findings (Adam et al., 2018, 2020; Medendorp et al., 2007; Sauseng et al., 2009; van der Werf et al., 2008; van Dijk et al., 2010), while the topography of alpha power can be used to decode the location of items held in WM (Bae & Luck, 2018; Foster et al., 2016). In contrast, theta’s role in WM is less clear. While rodent hippocampal theta oscillations strongly relate to behavior and cognition (Soltani Zangbar et al., 2020; Vanderwolf, 1969), the evidence for frontal midline theta in humans is sparser and mixed (Herweg et al., 2020; Pavlov & Kotchoubey, 2022).

The lack of clarity regarding the role of theta might be explained by non-oscillatory, aperiodic neural dynamics. Traditional time-frequency analysis methods conflate oscillations with aperiodic activity (Donoghue, Haller, Peterson, et al., 2020; He, 2014). Historically often treated as task-irrelevant noise, a growing consensus shows aperiodic activity is dynamic and task-relevant in aging (Voytek et al., 2015), sleep (Höhn et al., 2024; Lendner et al., 2020), anesthesia (Colombo et al., 2019; Waschke et al., 2021), and disease (Helson et al., 2023; Karalunas et al., 2022; Smith, Kosik, van Engen, et al., 2023; Smith, Ma, et al., 2023). Preliminary evidence suggests that resting-state aperiodic activity is correlated with WM capacity (McKeon et al., 2024), and task-based aperiodic activity is involved in WM processes (Frelih et al., 2024; Gyurkovics et al., 2022; Virtue-Griffiths et al., 2022).

*Here, we tested the hypothesis that the mixed results regarding human theta in WM can potentially be resolved by event-related aperiodic dynamics*. While it is possible the same applies to occipitoparietal alpha activity – that it might be driven partly by aperiodic activity - previous work suggests that alpha oscillations are less likely to be masked by aperiodic dynamics than theta (Donoghue, Dominguez, et al., 2020). Here we reanalyzed two (n = 74) human EEG datasets of participants performing a visual WM task (Fig. 1A) (Adam et al., 2018). Using spectral parameterization (Fig. 1B), we replicated traditional alpha effects when accounting for aperiodic activity: power decreased with increasing WM load. The original paper (Adam et al., 2018) used the filter-Hilbert method and reported task-related theta power decreases during WM retention. In contrast, we found a decrease in aperiodic activity, but not in aperiodic-adjusted theta power.

**Figure 1.**
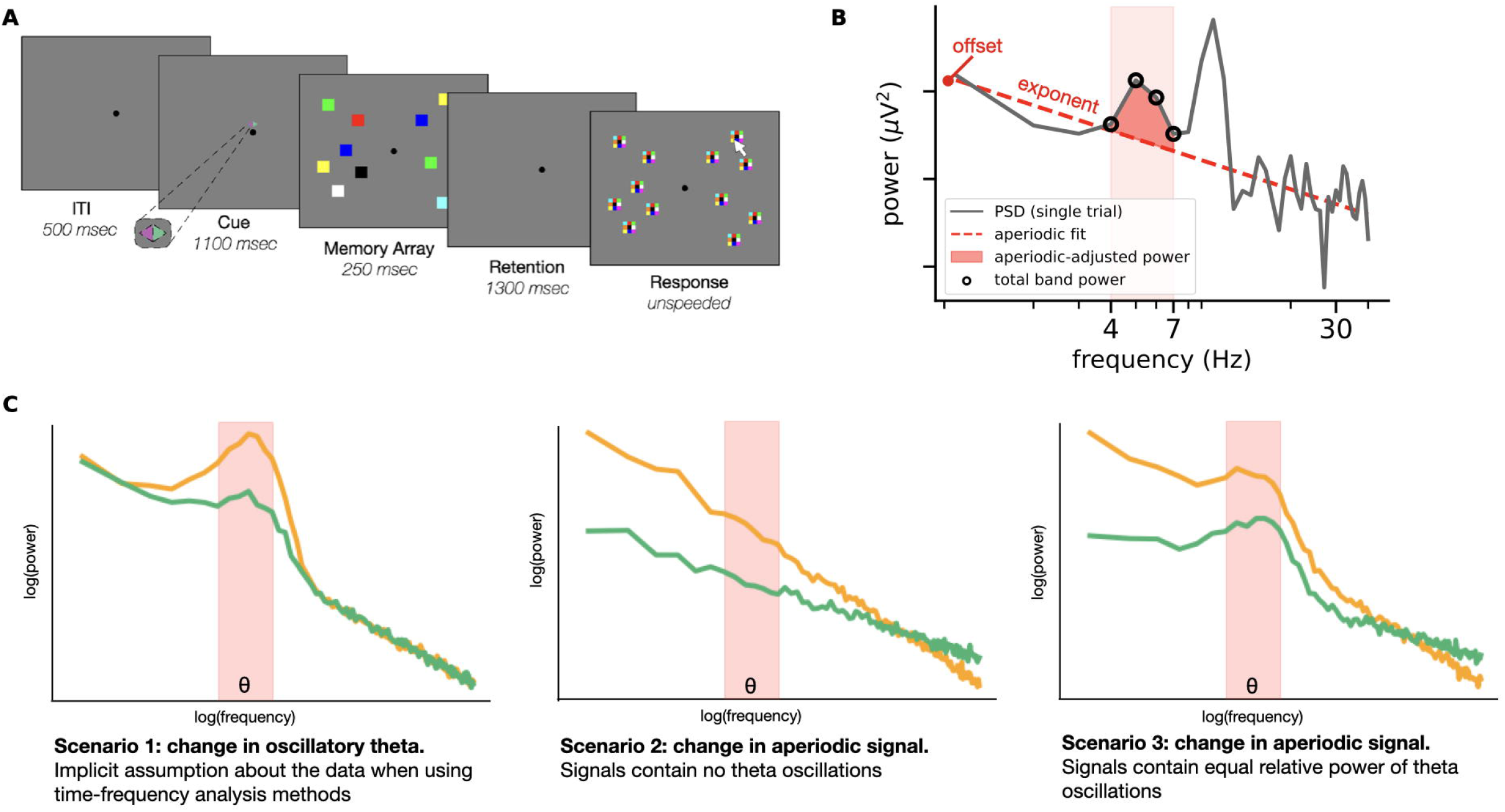
Overview of experiment, analysis, and hypotheses. A) Visual description of the whole-report visual working memory task (adapted from Adam et al., 2018). A cue indicated whether participants were required to attend to the left or right side of the screen. Both sides contained the same number of colored squares, but in different locations. In experiment 1, set size was manipulated to consist of one, three or six items on each side. In experiment 2, the set size was kept constant at six items. After a 1300 ms retention period, participants were prompted to indicate the color of each item of the attended side. There were nine discrete colors to choose from. B) Spectral parameterization method. From a power spectrum, the aperiodic components, the exponent and offset, were quantified separately from the periodic components, aperiodic-adjusted power. Total band power was captured as the average power within the frequency band of interest. C) Three scenarios illustrating how task-induced changes in the power spectrum appear as theta total band power changes. The theta frequency range (4-7 Hz) is highlighted in red. Scenario 1 (left) shows the assumed model whereby task-related changes in theta band activity are driven by changes in narrowband, oscillatory theta activity manifesting above a stable aperiodic signal. Scenario 2 (middle) and 3 (right) describe situations in which a reduction in total theta power on the task (green line) from baseline (orange line) is observed, all driven by changes in aperiodic dynamics whether detectable theta oscillations are present or not.

By disentangling aperiodic activity from narrowband oscillations, we found task-general changes in aperiodic activity (spectral flattening) in the frontal midline and occipitoparietal regions. We further found occipitoparietal task-dependent effects; aperiodic flattening with increased WM load. Consistent with prior research (Bailey et al., 2020; Klimesch, 1999; Mitchell et al., 2008), we found evidence for theta oscillations in fewer than half (42%) of the participants, whereas almost every participant (91%) showed clear alpha oscillations. Our results demonstrate how implicit assumptions of time-frequency analysis methods can lead to misleading conclusions. We show correlational evidence of aperiodic activity in human visual WM, demonstrating new insights into oscillatory dynamics by accounting for event-related aperiodic activity.

## 2. Methods

### 2.1. Study design from Adam et al. (2018)

All data used in this manuscript was obtained from OSF (https://osf.io/8xuk3/) (Adam et al., 2018).

#### 2.1.1. Participants

Separate groups of participants were recruited for Experiments 1 and 2. All participants were between the age of 18 and 35, with normal or corrected-to-normal visual acuity and color vision, and of either sex. Data from 31 participants were collected in Experiment 1, and 48 participants in Experiment 2. For more details on inclusion and exclusion criteria, see Adam et al. (2018). After EEG artifact rejection, 30 participants were considered for further analysis in Experiment 1, and 44 in Experiment 2.

#### 2.1.2 Task design

In both experiments, participants performed a whole-report visual WM task (Fig. 1A), in which they were asked to remember the colors of differently colored squares across a blank screen retention period and then recall the colors of all remembered items (Huang et al 2010; Adam et al, 2015). Throughout each trial, participants were asked to fixate on a dot presented in the middle of the screen, except during the response. A cue was presented for 1100 ms to indicate whether they had to memorize the items on the right or the left side of the screen. The memory array was presented for 250 ms followed by a retention period of 1300 ms, after which a response for each cued item had to be given (Fig. 1B). The number of items were equal on each side to balance the visual stimulation within each visual hemifield. This design was chosen to ensure hemispheric differences in neural activity were driven by cognitive demands, rather than perceptual demands (Vogel & Machizawa, 2004).

Each block consisted of 30 trials. For Experiment 1, each block had 10 trials for three different set sizes, one, three or six, and Experiment 2 contained 30 trials of set size six. The conditions in Experiment 1 were defined as the three set sizes, whereas in Experiment 2 two conditions were defined based on the performance on each trial: if more than three items were correctly answered, it was deemed a “high performance” trial, versus having less than three items correct was deemed a “poor performance” trial.

#### 2.1.3 EEG acquisition

EEG passive electrodes and cap were from ElectroCap International (Eaton, OH). Recordings were done on an SA Instruments Amplifier running LabView 6.1 (Fife, Scotland). The cap contains 20 electrodes according to the 10/20 international sites. Two supplementary electrodes were placed over the occipital lobe: OL was placed between T5 and O1, and OR was placed between T6 and O2. In addition, electrooculography (EOG) was recorded to measure horizontal eye movements and blinks. Data were band-pass filtered between 0.01-80 Hz with a sampling frequency of 250 Hz. Further information is available in previous reports on this dataset (Adam et al., 2018; Foster et al., 2016).

#### 2.1.4 EEG preprocessing

Data were available from the original paper’s repository, already pre-processed and epoched, but not baselined. Pre-processing steps included: automatic removal of trials with eye blinks (window size of 200 ms, step size of 10 ms, and a threshold of 50 μV) or eye movements (split-half sliding window of size 200 ms, step size of 10 ms, and a threshold of 20 μV), automatic removal of trials with excessive noise using a peak-to-peak threshold rejection (a change greater than 200 μV within a 15 ms window), and finally a visual inspection of the data to determine whether the automated steps were successfully implemented. More details can be found in Adam et al. (2018). Here, we used the non-baselined epochs.

The same electrode clusters were used as defined by the original paper (Adam et al., 2018) for replicating their results, and the novel analysis conducted in this paper. These clusters for lateralized alpha were left and right occipital and parietal regions (O1, OL, P3, PO3, T5, O2, OR, P4, PO4, and T6), and for theta a cluster over the frontal-midline (F3, F4, Fz).

### 2.2 Replication of original results

To replicate the original results (Adam et al., 2018), we performed the same analysis. We separate bandpass filtering in the theta (4-7Hz) and alpha (8-12Hz) bands. We then performed a Hilbert transform to compute the analytic amplitude of these frequency bands over time. This amplitude was baseline corrected to 400 ms before cue onset.

For statistical analysis, theta and alpha power was averaged over the retention period (400 - 1500 ms after memory array onset) per condition. A two-way repeated-measures ANOVA was performed on alpha power with condition (set size for Experiment 1 or performance for Experiment 2) and lateralization as factors. Note that the factor set size or performance is comparable to a whole-field alpha approach, and the second factor adds the lateralization effect. A pairwise t-test was used on theta power to investigate the difference between good and poor performance.

### 2.3 Spectral parameterization: separating oscillatory from aperiodic activity

#### 2.3.1 The specparam model

Spectral parameterization (Donoghue, Haller, Peterson, et al., 2020; https://specparam-tools.github.io/) models a power spectrum as a combination of an aperiodic component (fit as a Lorentzian) and putative oscillatory components that rise above the aperiodic signal (fit as Gaussians). Here, the parameters characterizing the aperiodic component are the spectral exponent and the offset. Three parameters characterize the Gaussian periodic components: peak height (relative to the aperiodic signal), center frequency, and bandwidth (Fig. 1B). More details can be found in (Donoghue, Haller, Peterson, et al., 2020).

#### 2.3.2 Identification of participants exhibiting oscillatory activity

Because oscillations are not consistently present (Jones, 2016), spectral parameterization helps to explicitly identify participants and electrodes where there are putative oscillations (defined here as having a significant “bump” in the power spectrum within the theta or alpha range above aperiodic activity). This triggered an additional inclusion criterion, different from the original report (Adam et al., 2018), in that oscillatory activity should only be analyzed from participants who exhibited putative oscillations. Thus, for alpha power in Experiments 1 and 2, participants were required to have a parameterized spectral peak between 8 and 12 Hz in the occipitoparietal electrodes. For theta power, participants were required to have a peak between 4 and 7 Hz that was distinct from alpha activity on frontal midline electrodes (detailed below). To perform this analysis, a power spectrum was generated for every trial, for the whole trial length using Welch’s method and a Hanning window of two seconds without overlap. The model was applied to the power spectrum in the frequency range of 2-40 Hz. Peak width limits were set from 2-8 Hz. To minimize overfitting of relatively flat spectra without peaks, *specparam* requires an amplitude threshold for determining the height of the Gaussian, relative to the aperiodic signal, to be considered a peak. This threshold was set at 0.2 µV^2^/Hz for the alpha frequency band, and 0.1 µV^2^/Hz for theta frequency band. Different absolute thresholds were chosen for the bands because theta power is typically lower than alpha power, relative to the aperiodic fit. For both experiments, *specparam* successfully discriminated between participants who exhibited alpha power and those who did not. This was further confirmed by visual inspection. However, discriminating participants based on theta power proved to be more difficult because not all participants exhibited a distinct theta peak. A portion of participants also exhibited alpha power on the frontal midline electrodes overlapping with the theta frequency range. Therefore, determining whether a participant exhibits theta power was guided by *specparam* and confirmed with manual visual classification to avoid lower frequency alpha peaks to be considered theta.

#### 2.3.3 Generating single trial power spectra and applying specparam

Power spectra were created for each trial, each window of interest (cue and retention period), and for each electrode of interest. Then, spectra were averaged over electrodes within each electrode group, contralateral occipitoparietal, ipsilateral occipitoparietal, and frontal midline. A time window from 400 to 1500 ms after memory array onset (1100 ms) was used for the retention period to produce power spectra, excluding the early perceptual ERP time points (Kałamała et al., 2024). For a baseline, we used the cue period from −1100 until the memory array onset (1100 ms). This decision is due to the ITI only being 500 ms long, which does not provide a sufficient baseline otherwise. Power spectra were created using the Fast Fourier Transform with a Hanning window equal to the length of the time windows of (both cue and retention windows are 1100 ms). The *specparam* model was applied to these power spectra using the same settings described in section 2.2.2. Five outputs were collected per trial (Fig. 1B):

1. Aperiodic-adjusted oscillatory (alpha or theta) power was calculated by taking the area under the curve (AUC) between the power spectrum and aperiodic fit on the logarithmic scale. The AUC was calculated between 4-7 Hz for theta, and 8-12 Hz for alpha (Bender et al., 2025; Cunningham et al., 2023).
2. Abundance of oscillations, defined as the percentage of trials within a condition in which a peak in the alpha or theta range is observed.
3. The aperiodic exponent.
4. The aperiodic offset.
5. Total band power was calculated as the average power between 4-7 Hz for theta, and 8-12 Hz for alpha.

A baseline correction was performed by subtracting the values of these parameters during the cue period from the values of the retention period in a trial-by-trial fashion for all outputs (except abundance) as is common for task-related spectral parameterization analyses (Donoghue, Haller, et al., 2020). Afterwards, each output was averaged for each condition and participant.

#### 2.3.4 Inclusion criteria for single-trial model fit

Model fit inspection was done by calculating the mean R^2^ per condition for each participant. All trials with a model fit lower than two times the standard deviation of the mean were removed from further analysis. On average, 8% of trials were removed for each condition per participant. Afterwards, participants were checked again ensuring they have a minimum number of trials per condition (75 in Experiment 1, and 40 in Experiment 2) to remain included in this study. This resulted in excluding four participants in Experiment 1 and another four in Experiment 2 for the occipitoparietal analysis. No participants were excluded based on this criterion in either experiment for the frontal midline analysis. The average model fit and standard deviations for the included participants in Experiment 1 were 0.71±0.09 for occipitoparietal and 0.70±0.09 for frontal midline analyses. For Experiment 2, they were 0.69±0.09 for occipitoparietal, and 0.66±0.09 for frontal midline analyses. Note that due to the short amount of time per window, these model fits are lower than what is typically seen when parameterizing data that is averaged across trials and/or over long segments of resting-state EEG. For our analysis design, we required measures of individual trials, thereby limiting the amount of data available per model fit.

We observe statistically significant differences in model fits between the cue and retention period for the occipitoparietal clusters in both Experiments, and for the frontal-midline cluster in Experiment 1 (p<0.01). The cue period had worse fit than the retention period on occipitoparietal electrodes, but this direction was reserved for the frontal-midline. Furthermore, the ipsilateral hemisphere had significantly higher model fits than the contralateral hemisphere for both Experiments (p<0.01). In Experiment 1, model fits were significantly decreased with an increase in set size for the occipitoparietal clusters (p<0.001), but not for the frontal-midline. In Experiment 2, we observed the reversed effect; model fits were significantly higher for the good performance, than the poor performance (p<0.05). Even though differences were significant, we consider them negligible since the largest observed distance between averages was only 0.01.

### 2.4 Behavioral analysis and correlation to EEG metrics

WM capacity was calculated in two ways. The first was based on the main task while EEG was recorded, by calculating the average number of items correctly remembered over all trials. The second method was based on the performance during a baseline WM task which followed the same procedure in both experiments (referred to as the Color Change Detection Task in Adam et al., (2018). This WM estimate was calculated per set size, and then averaged to get a single value representing WM capacity per participant, expressed as an estimate (“K”) which is calculated with the following formula:

K = N x (H - FA)

N = set size

H = hit rate (proportion of correct trials)

FA = false alarm rate (proportion of incorrect no-change trials)

### 2.5 Statistical analysis

The dependent variables in both experiments were the oscillatory power relative to the aperiodic signal, oscillation abundance, aperiodic exponent, aperiodic offset, and total band power.

For the occipitoparietal region, separate two-factor repeated measures ANOVAs were applied on all dependent variables. The factors were set size in Experiment 1 (one, three, six) or performance in Experiment 2 (good, poor) and lateralization (ipsilateral, contralateral). In cases where the ANOVA was significant, pairwise t-tests with Bonferroni correction were applied as post-hoc tests to examine the differences between the levels within each factor. For the frontal midline region, separate repeated measures ANOVA were applied to test for significant differences on all dependent variables between set sizes (Experiment 1), and pairwise t-tests were applied similarly to test statistical significance of performance (Experiment 2). Additionally, paired t-tests were applied on each variable averaged over the conditions to test whether there was a significant difference between the retention and cue period.

Shapiro-Wilk tests were used to test for normality, Levene’s test for equal variance, and Mauchly’s test were used to test for sphericity. If sphericity was violated, the Greenhouse-Geisser correction was applied for the factorial repeated measures ANOVAs. If one or more assumptions were violated, a Friedman test was used instead of a repeated measures ANOVA, or the Wilcoxon signed rank test was used instead of a pairwise t-test. Cohen’s d was reported as effect size for t-tests, partial eta-squared for ANOVA tests, and the Common-Language Effect Size (CLES) for non-parametric tests.

For our exploratory analysis, we investigated correlations between WM capacity and aperiodic exponent and offset. Two versions of WM capacity were captured, from a baseline task, and the main task. Furthermore, the pre-stimulus (cue period), retention period, and the change between these two windows were used. Lastly, there were two regions of interest, the frontal midline, and occipitoparietal regions. Thus, there were 24 correlations established using the Pearson correlations. To avoid finding spurious significance values, we applied a Bonferroni thresholding correction (α = 2.08*^10-3^).

### 2.6 Software

All data were processed using the Python programming language (version 3.7.1). Signal processing was done with the *neurodsp* module (version 2.1.0) (Cole et al., 2019), and power spectra were fitted with the *specparam* toolbox (version 1.0.0) (Donoghue, Haller, Peterson, et al., 2020). All statistics were performed in Pingouin (version 0.5.2) (Vallat, 2018), and plots were created using Matplotlib (version 3.0.2) (Hunter, 2007) and seaborn (version 0.9.0) (Waskom, 2021).

### 2.7 Data and code statement

Data used in this project is from Adam et al. (2018), and is available at OSF (https://osf.io/8xuk3/). The code for the both the replication, and analyses carried out in this paper are available on our GitHub page (https://github.com/voytekresearch/WM_specparam).

## 3. Results

We illustrate three scenarios where task-induced changes to neural power spectra appear as theta total band power decreases (Fig. 1C). The first illustrates the - potentially incorrect - standard model, whereby aperiodic activity is stable between conditions while narrowband theta oscillatory power dynamically changes. The second and third scenarios show no difference in aperiodic-adjusted theta power (area of the bump), whereas total theta band power does. Here, theta band effects are driven by aperiodic dynamics, regardless of whether detectable theta oscillations are present.

### 3.1 Successful replication of the original paper’s results

All analyses from the original paper (Adam et al., 2018) were successfully replicated before moving onto the spectral parameterization analysis. These results can be found on our GitHub repository. In short, in Experiment 1 we observed larger differences between hemispheres in occipitoparietal alpha power with an increase in WM set size. In Experiment 2, when separating conditions by their accuracy, theta power was higher for good WM performance than poor.

### 3.2 Spectral parameterization: separating oscillatory from aperiodic activity

#### 3.2.1. Participants

For this analysis, only participants exhibiting alpha oscillations over occipitoparietal electrodes or theta power over frontal midline electrodes were included. Furthermore, they needed to pass the minimum number of trial requirements as well. All 30 participants in Experiment 1, and the majority (37 out of 44) in Experiment 2 exhibited clear peaks in the alpha range according to the spectral parameterization criteria (Fig. 2). After excluding participants who did not meet the minimum number of trials requirement, we ended with a sample size of 26 in Experiment 1, and 26 in Experiment 2. In contrast, only 12 out of 30 participants in Experiment 1, and 19 out of 44 in Experiment 2 met the criteria for exhibiting putative theta oscillations, identified as peaks in the theta range (Fig. 2). The final sample size after excluding participants is 10 in Experiment 1, and 17 in Experiment 2.

**Figure 2.**
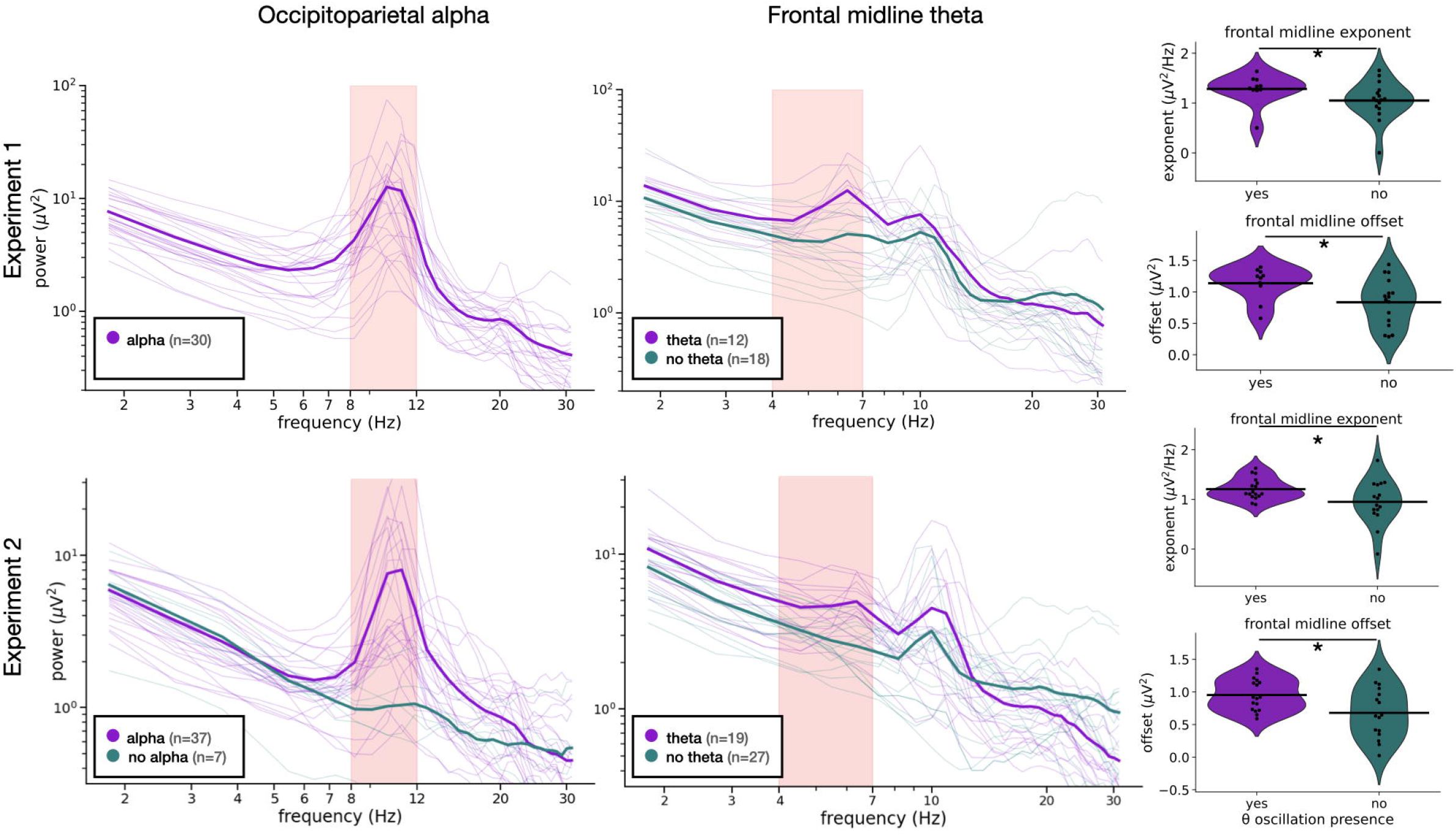
Grouping participants based on the presence of oscillatory peaks. Here we show for both experiments (rows), and oscillatory bands (columns) the average (thick lines), and individual participant PSDs (thin lines) averaged over all trials for those who did exhibit oscillatory power (magenta), and those who did not (teal). For theta, we also observe group differences in both aperiodic exponent and offset, in which participants without theta oscillations typically have flatter spectra. These results apply to both experiments. * = p < 0.05

To increase the sample size and therefore statistical power for the aperiodic analyses on frontal midline electrodes, we decided to include all participants, exhibiting putative theta oscillations or not. In Experiment 1, 16 from the 18 participants without oscillations satisfied the minimum trial count requirement and were included here, bringing the total to 26 participants. In Experiment 2, 16 from the 27 met these criteria in Experiment 2, bringing that total to 33.

#### 3.2.2 Experiment 1: Frontal midline aperiodic-adjusted theta power and aperiodic activity

From the 12 participants with putative theta oscillations, 2 were rejected due to low trial count. Power spectra are shown for the remaining 10 participants included in this analysis concerning theta oscillations (Fig. 3A), aperiodic-adjusted theta power was not significantly different between set sizes (Q = 2.6, p = 0.27, W = 0.13), nor was it significantly increased from the cue period over the three set sizes was significantly (t(9) = 1.69, p = 0.13, Cohen’s d = 0.40; Fig. 3B). Like the aperiodic-adjusted theta power results, theta abundance was not significantly different between set sizes (Q = 0.2, p = 0.90, W = 0.01). In contrast to power, there was a significant decrease in abundance in the retention period from the cue period over all conditions (W = 7, p = 0.037, CLES = 0.36; Fig. 3D).

**Figure 3.**
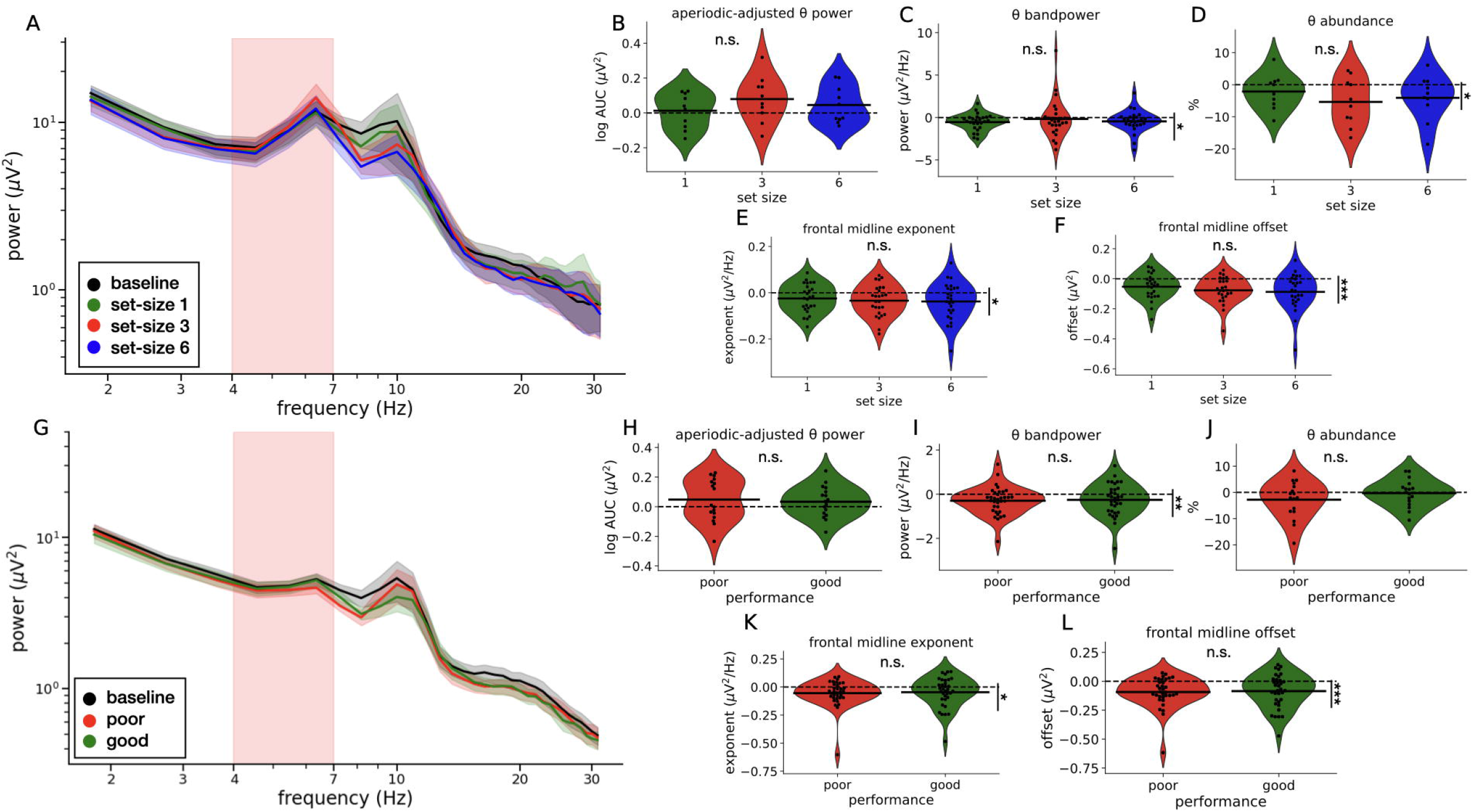
Frontal midline aperiodic activity decreases during the retention period from the cue period. A-F) Experiment 1 results. A) Shows the power spectra from baseline (black), and the three different set sizes.B) Aperiodic-adjusted theta power as the logarithmic AUC between the PSD and aperiodic fit is not significantly different between set sizes. Theta abundance (D) and total band power (C) both decrease during the retention period from the cue period. Aperiodic exponent (E) and offset (F) show no significant differences between set sizes, but significantly decreased from baseline. G-L) Experiment 2 results. F) Shows the power spectra for the baseline (black), and poor (red) versus good (green) WM performance. No significant differences were observed in aperiodic-adjusted theta power (H), nor in abundance (J), whereas both total band power (I), aperiodic exponent (K), and offset (L) decreased from the cue period to the retention period. * = p < 0.05, ** = p < 0.005, ***= p < 0.001, n.s. = not significant.

Frontal midline aperiodic exponent and offset were calculated for all participants included (n = 25), not just the participants that also had theta power. The frontal aperiodic exponent was not significantly different between set sizes (Q = 2.15, p = 0.34, W = 0.04), but was significantly decreased from the cue period over all conditions (W = 66, p = 5.6*10^-3^, CLES = 0.48; Fig. 3E). A similar pattern was observed for the frontal midline offset; no significant differences between set sizes (Q = 2.38, p = 0.30, W = 0.046), but aperiodic offset was significantly reduced from the cue period over all conditions (W = 0.046, p = 8.3*10^-5^, CLES = 0.44; Fig. 3F). Theta total band power was not significantly different between set sizes (Q = 1.46, p = 0.48, W = 0.028), but was also significantly reduced during the retention period over all set sizes (W = 89, p = 2.9*10^−2^, CLES = 0.46; Fig. 3C).

These results, leveraging the novel spectral parameterization method, show that aperiodic activity (exponent and offset) is significantly reduced during the WM maintenance period from our baseline over frontal midline regions, but no differences were observed between WM loads. Furthermore, data suggests that theta oscillations are less frequently observed in the retention period compared to the cue period, but we found no differences in theta power between WM loads. Notably, the original paper (Adam et al., 2018) did not report frontal midline theta power results for Experiment 1. We have implemented this analysis, and did not observe WM load effects on frontal midline theta power using the Hilbert transform approach (see our GitHub repository for statistics and the figure). Thus, our alternative aperiodic-adjusted theta power results are in line with the original methodological approach.

#### 3.2.3 Experiment 2: Frontal midline aperiodic-adjusted theta power and aperiodic activity

Seventeen participants met all inclusion criteria for theta oscillation analyses, and their power spectra are shown in Fig. 3G. Aperiodic-adjusted theta power was not significantly different between poor and good performance (t(16) = −0.44, p = 0.67, Cohen’s d = −0.12), nor was it different during the retention period from the cue period (t(16) = 1.64, p = 0.12, Cohen’s d = 0.29; Fig. 3H). Similarly, theta abundance was not significantly different between poor and good performance (t(16) = 1.08, p = 0.30, Cohen’s d = 0.41), nor was it different between the cue and retention period (t(16) = −1.62, p = 0.12, Cohen’s d = 0.23; Fig. 3J).

The frontal midline aperiodic exponent and offset were calculated for all included participants (n=34), regardless of exhibiting putative theta oscillations or not. Neither of the aperiodic measures were normally distributed; therefore, the non-parametric Wilcoxon signed-rank test was used to determine statistical significance. The exponent was not significantly different between performances (W = 280, p = 0.77, CLES = 0.54), but it was significantly decreased from the cue period (t(32) = −2.37, p = 2.4*10^-2^, Cohen’s d =0.14; Fig. 3K). Aperiodic offset was not significantly different between performances (W = 283, p = 0.81, CLES = 0.52), offset was significantly reduced from the cue period (t(32) = −4.09, p = 2.7*10^-4^, Cohen’s d = 0.27; Fig. 3L). Theta total band power showed no difference between performances (t(25) < 1.0), but was significantly reduced during the retention period from the cue period (W = 135, p = 9.6*10^-3^, CLES = 0.45; Fig. 3I).

Similar to Experiment 1, task state caused a significant reduction in aperiodic exponent and offset, with no differences between poor and good performance. In contrast to Experiment 1, we did not find differences in theta abundance by task-state (cue vs. retention) nor by task performance (poor vs. good). Aperiodic-adjusted theta power was not significantly higher in the good performance condition compared to poor, as we expected based on the original paper (Adam et al., 2018). Thus, with spectral parameterization, we were unable to replicate the original theta power differences between good and poor WM performance with either aperiodic exponent or offset, nor with aperiodic-adjusted theta power.

#### 3.2.4 Experiment 1: Occipitoparietal aperiodic-adjusted alpha power and aperiodic activity

Power spectra for the twenty-six included participants are shown in Fig. 4A. Using a two-way repeated measures ANOVA, there was a main effect of set size on aperiodic-adjusted occipitoparietal alpha power (F(2,50) = 6.55, p = 6.0*10^-3^, η _p_^2^ = 0.20), but not for lateralization (F(1,25) = 0.62, p = 0.43, η _p_^2^ = 0.024) or the interaction between set size and lateralization (F(2,50) = 2.95, p = 6.1*10^-2^, η _p_^2^= 0.11). Alpha power in set size one was significantly higher than set size six (t(25) = 3.27, p = 3.1*10^-3^, Cohen’s d = 0.36), and between set size three and six (t(25) = 2.20, p = 3.7*10^-2^, Cohen’s d = 0.24), but not different between set size one and six (t(25) = 1.82, p = 8.1*10^-2^, Cohen’s d = 0.36; Fig. 4B). Alpha abundance was significantly different between set sizes (F(2,50) 3.59, p = 3.5*10^-2^, η _p_^2^ = 0.13), with the post-hoc only showing a significant increase from set size one to six (t(25) = −2.44, p = 2.2*10^-2^, Cohen’s d = −0.40; Fig. 4C). Main effect of lateralization was not significant, nor in the interaction (F < 1.0).

**Figure 4.**
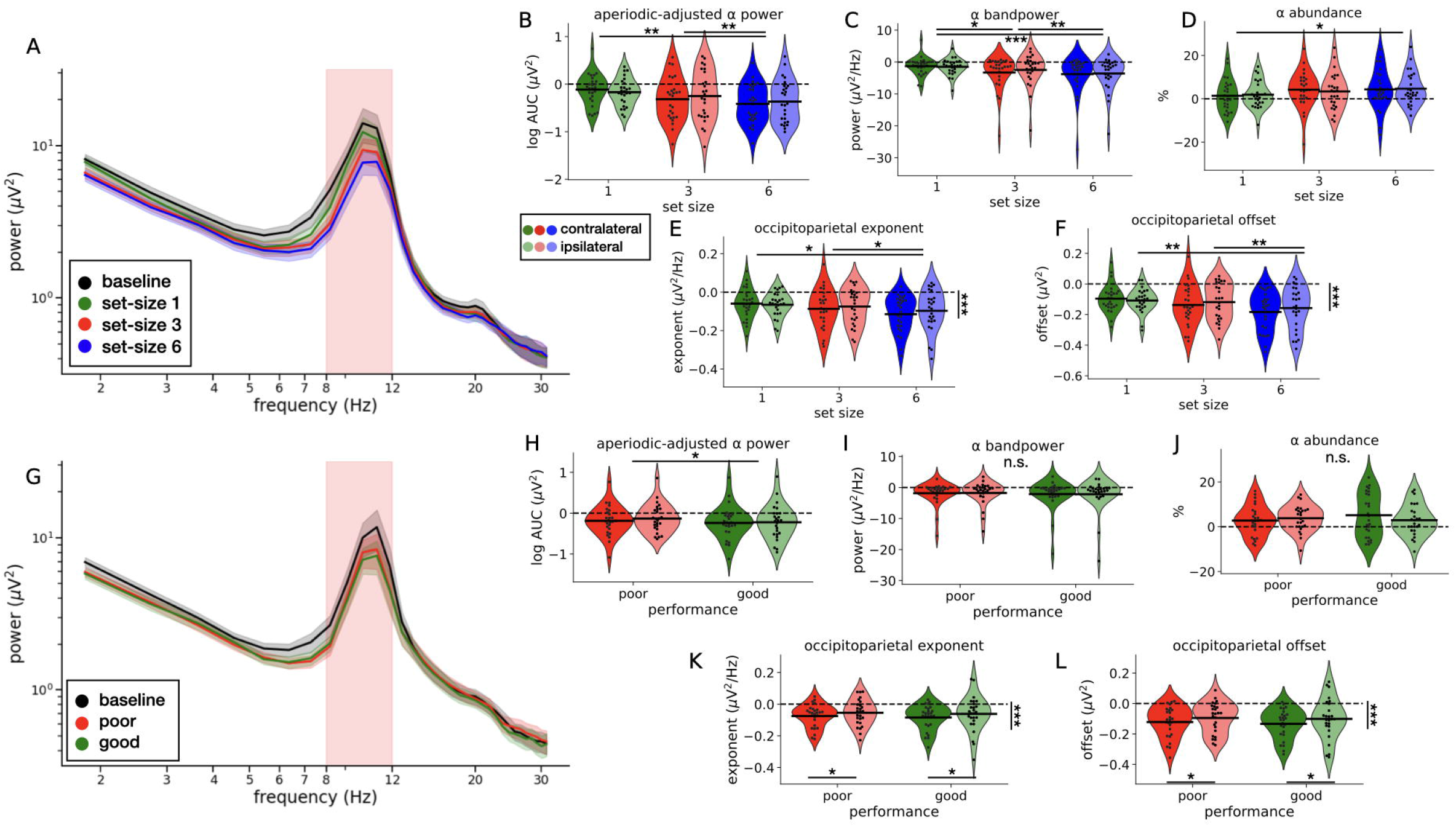
Occipitoparietal aperiodic activity decreases with increase in load, but not performance, whereas alpha oscillatory power follows classic findings. A-F) Experiment 1 results. A) Shows the power spectra from baseline (black), and three set sizes. B) Aperiodic-adjusted occipitoparietal alpha power as the logarithmic AUC between the PSD and aperiodic fit showed a significant decrease with an increase in WM load, but no lateralized effect is observed. Similar patterns are observed for total alpha total band power (C), whereas abundance increases with set size (D). Aperiodic exponent (E) shows no lateralized effect, but there was a decrease with an increase in WM load. Similar patterns were observed for the aperiodic offset (F). G-L) Experiment 2 results. F) Shows the power spectra for the baseline (black), and poor (red) versus good (green) WM performance. A decrease in aperiodic-adjusted occipitoparietal alpha power was observed for the good condition compared to the poor condition, but again no lateralization effect was found (H). Total band power and (I) abundance (J) did not show any significant differences. The aperiodic exponent (K) and offset (L) were both significantly reduced from baseline, and a lateralization effect was observed, in which activity in the contralateral hemisphere was more suppressed than the ipsilateral hemisphere. * = p < 0.05, ** = p < 0.005, ***= p < 0.001, n.s. = not significant

There was a main effect of set size on occipitoparietal aperiodic exponent such that it decreased with increasing set size (F(2,50) = 5.99, p = 4.6*10^-3^, η _p_^2^= 0.19). A post-hoc test shows that the difference is significant between set sizes one and six (t(25) = 3.08, p = 5.0*10^-3^, Cohen’s d = 0.52), and between set sizes three and six (t(25) = 2.35, p = 2.7*10^-2^, Cohen’s d = 0.26; Fig. 4E). No significant effect was found for lateralization, nor was there an interaction effect between lateralization and set size (F < 1.0). Furthermore, exponent was significantly lower during the retention period compared to the cue period over all conditions (t(25) = −5.43, p = 1.2*10^-5^, Cohen’s d = 0.22). Occipitoparietal aperiodic offset was also significantly decreased with increasing set sizes (F(2,50) = 7.68, p = 1.2*10^-3^, η _p_^2^ = 0.23). A post-hoc pairwise t-test showed significant differences between set sizes one and six (t(25) = 3.30, p = 2.9*10^-3^, Cohen’s d = 0.64), and between set sizes three and six (t(25) = 3.01, p = 5.9*10^-3^, Cohen’s d = 0.34; Fig 4F). Furthermore, over all conditions the offset was also significantly lower during the retention period compared to the cue period (t(25) = −6.93, p = 2.9*10^-7^, Cohen’s d = 0.32). Total band power showed a main effect of set size (F(2,50) = 7.18, p = 1.8*10^-3^, η _p_^2^ = 0.22), but not for lateralization (F(1,25) < 1.0) or their interaction (F(2,50) = 2.98, p = 6.0*10^-2^, η _p_^2^= 0.11). Post-hoc tests showed that all three set sizes were significantly different from each other. Set size one had the lowest total band power, lower than set size three (t(25) = 2.15, p = 4.2*10^-2^, Cohen’s d = 0.36), and set size six (t(25) = 2.95, p = 6.8*10^-3^, Cohen’s d = 0.54), and set size three was lower than set size six (t(25) = 3.74, p = 9.5*10^-4^, Cohen’s d = 0.15; Fig. 4C).

In summary, we observed aperiodic-adjusted alpha power decreases with an increase in WM set size, but no differences between hemispheres. A lack of lateralization is in contrast with the original paper’s findings (Adam et al., 2018). However, when we look at the data without subtracting the cue period as a baselining method, we observe similar lateralization effects (F(1,25) = 22.81, p = 6.7*10^-5^, η _p_^2^ = 0.48), including a strong interaction between set size and lateralization (F(2,50) = 6.82, p = 2.4*10^-3^, η _p_^2^ = 0.21). Adam et al. (2018) also reported finding strong lateralization effects during the cue period. Hence, these effects are cancelled out when we use the cue period as our baseline. The occipitoparietal aperiodic exponent and offset showed a decrease with an increase in WM set size, on top of an overall decrease during the retention period from the cue period. Thus, we have replicated the original results with aperiodic-adjusted alpha power, in addition to finding that aperiodic activity is involved in WM processes over occipitoparietal regions.

#### 3.2.5 Experiment 2: Occipitoparietal aperiodic-adjusted alpha power and aperiodic activity

Power spectra are shown for the twenty-six included participants in Fig. 4G. For aperiodic-adjusted alpha oscillations, we observed a significant main effect of performance (F(1,25) = 4.25, p = 4.9*10^-2^, η _p_^2^ = 0.15; Fig. 4H), in which power was lower in the good performance condition compared to the poor performance condition. However, neither the lateralization nor the interaction effect were significant (F < 1.0). Alpha abundance did not differ as a function of performance, lateralization, or their interaction (F < 1.0; Fig. 4J).

Occipitoparietal aperiodic exponent was not significantly different between performance, nor the interaction between performance and lateralization (F < 1.0), but it was for lateralization (F(1,25) = 4.34, p= 4.8*10^-2^, η _p_^2^= 0.15; Fig. 4K) in which the contralateral hemisphere had a lower exponent than the ipsilateral hemisphere. The exponent was also significantly reduced during the retention period compared to the cue period over all conditions (t(25) = −4.99, p = 3.9*10^-5^, Cohen’s d = 0.24). A similar pattern was observed for the offset; the contralateral hemisphere had a lower offset than the ipsilateral hemisphere (F(1,25) = 6.08, p = 2.1*10^-2^, η _p_^2^, = 0.20; Fig. 4L), and an overall reduction was found compared to the cue period (t(25) = −6.41, p = 1.0*10^-6^, Cohen’s d = 0.38). Whereas alpha total band power showed a main effect of set size in Experiment 1, no significant results were observed between good or poor performance in Experiment 2 (F < 1.0; Fig. 4I).

Aperiodic-adjusted alpha power was lower on good performance trials than poor, which is in line with our results for set sizes in Experiment 1. The lack of lateralization effect observed here is comparable to Experiment 1; when considering the retention window without subtracting the cue period as a baselining method, we do observe a strong lateralization effect of aperiodic-adjusted alpha power (F(1,25) = 27.9, p = 1.8*10^-5^, η _p_^2^= 0.53). Aperiodic activity (exponent and offset) was lower in the contralateral hemisphere, compared to the ipsilateral hemisphere, but no effect of performance was found. In addition, we also observed an overall reduction in aperiodic exponent and offset during the retention period from the cue. Thereby, we both datasets reanalyzed here support our hypothesis that aperiodic activity is functionally involved in WM.

### 3.3 Correlations between working memory capacity and task-states

Given the functional role of theta oscillations in WM, we next sought to examine whether there was a difference in WM capacity between participants for whom there was, versus was no detectable theta oscillations with spectral parameterization. To test this, we looked at WM capacity in two ways, either as estimated from the performance on the WM task performed during the EEG experiment, or as measured from a baseline WM task collected prior to EEG recording. Notably, these two metrics of WM capacity were correlated within participants for both Experiment 1 (r(25) = 0.56, p = 3.37 * 10^-3^) and Experiment 2 (r(33) = 0.41, p = 1.8*10^-2^).

We found no significant WM capacity differences between the groups of participants who exhibited or did not exhibit theta oscillations when measured from the main EEG task or from the baseline task, in either Experiment (p > 0.4 in all instances). This indicates that WM capacity is not reliant on whether participants exhibited measurable theta oscillations.

In an additional exploratory analysis, we examined whether either measure of WM capacity correlated to aperiodic activity, exponent or offset from frontal midline or occipitoparietal regions. We examined this using average pre-stimulus (cue period) aperiodic activity, during the retention period, and in the within-participant change from cue to retention. This resulted in 24 correlations, with a Bonferroni corrected threshold of p < 2.08*10^-3^. None of those correlations were significant under this criterion.

Finally, we noticed that there appeared to be a difference in the overall aperiodic-adjusted theta power between Experiments 1 and 2 (Fig. 3 A&G). Notably, in Experiment 1 set size could vary between one, three or six items, whereas in Experiment 2 the set size was constant at six items. To verify this visual observation, we performed t-test on relative theta power during the cue, retention period, and their difference between Experiment 1 and 2. Relative aperiodic-adjusted theta power was significantly higher during Experiment 1 for both the cue (t(15.85) = 2.57, p = 2.0*10^-2^, Cohen’s d = 1.09) and retention periods (t(12.49) = 2.56, p = 2.4*10^-2^, Cohen’s d = 1.17). However, the difference in theta oscillatory power between them was not significant (t(14.83) = 1.36, p =0.19, Cohen’s d = 0.59).

These results indicate that aperiodic-adjusted theta power is overall higher during Experiment 1 with a variable WM set size, compared to Experiment 2 with a constant high WM set size, but the task-induced changes in aperiodic-adjusted theta power from the cue period to the retention period are similar. More information and figures can be found on our GitHub page.

## 4. Discussion

Many theories of human visual WM rely on oscillatory mechanisms for coordinating information between brain regions, and as a neural code for memory (Lisman & Jensen, 2013; Palva & Palva, 2007). These theories are built on the assumption that event-related changes in those frequency bands always capture oscillations. We examined whether and to what degree theta and alpha oscillatory activity is distinct from aperiodic activity in WM. Explicitly controlling for aperiodic activity, we find that fewer than half of participants (42%) had an identifiable theta spectral peak. While not conclusive evidence for a lack of theta oscillations, this suggests that if theta oscillations are present in human WM, they are difficult to detect above aperiodic activity. Therefore, previous studies likely investigating theta’s role in WM may often capture non-oscillatory aperiodic activity in most participants, rather than true theta oscillations (Fig. 1c).

We showed aperiodic activity decreased during the WM retention period, whereas aperiodic-adjusted theta power non-significantly increased. This paradox, whereby theta simultaneously increases and decreases during WM retention, is mirrored by the mixed results regarding theta in human memory (Herweg et al., 2020; Pavlov & Kotchoubey, 2022). We believe this paradox can be explained by aperiodic dynamics, given that we observed significant aperiodic decreases during WM retention, but not between WM loads. If aperiodic dynamics are ignored, they can mask and muddle the underlying oscillatory theta effects. *Thus, mixed theta findings in human WM can partly be resolved by accounting for aperiodic dynamics*. The rarity of theta oscillations is an interesting contrast to the relative ubiquity of alpha oscillations. Even when accounting for aperiodic activity, we replicated the common finding that alpha power decreases with increasing WM load. A load-dependent decrease was also found for aperiodic activity. Thus, it is possible to observe simultaneous oscillatory and aperiodic task-related changes (Fig.1c), and we found both support visual attention and WM retention.

Importantly, we identified a task-general, load-dependent decrease of aperiodic activity during WM retention. Aperiodic activity has links to underlying physiological mechanisms, such as evidence that it captures the balance of excitation and inhibition (E:I) in a simple computational model (Gao et al., 2017), in pharmacological interventions (Colombo et al., 2019; Waschke et al., 2021), and using magnetic resonance spectroscopy imaging (McKeon et al., 2024). However, biophysical modeling work challenges this perhaps overly simplistic hypothesis (Brake et al., 2024). Nevertheless, if aperiodic activity serves as an indicator of E:I, we can make insightful connections with computational WM models that highlight the importance of a balanced E:I network to sustain activity for maintaining information (Lu et al., 2023; Wang, 1999). Thus, the task-general flattening of aperiodic activity across the brain could indicate a shift in E:I balance that is supported by computational models of sustained activity during the maintenance period. The task-specific modulations in occipitoparietal regions support the sensory recruitment theory, in which sensory regions are actively involved in the maintenance of WM items (D’Esposito & Postle, 2015). In this case, occipitoparietal regions help maintain visual stimuli through asynchronous and oscillatory activity, whereas activity from the frontal midline could be considered a top-down influence (Lara & Wallis, 2015).

While we found aperiodic flattening after stimulus onset, others found steepening or mixed results (Akbarian et al., 2023; Frelih et al., 2024; Kałamała et al., 2024; Lendner et al., 2023; Virtue-Griffiths et al., 2022). Notably, some of these papers (Akbarian et al., 2023; Frelih et al., 2024) used the n-back task, which mixes motor preparation responses with WM maintenance activity (Pavlov & Kotchoubey, 2022).

Mixed results were also observed on a visual continuous recall WM task; increased exponents over the frontal midline, and decreases over occipitoparietal regions, with an increase in exponent being associated with improved performance on a battery test (Virtue-Griffiths et al., 2022). Non-WM cognitive tasks suffer a similar fate; some find decreases in exponent after stimulus onset (Podvalny et al., 2015; Waschke et al., 2021), others find biphasic responses (Kałamała et al., 2024). These inconsistent results may be partly explained by the task and whether fast motor responses are required, but also potentially relate to differences in how aperiodic activity is measured and reported (Donoghue, 2024).

After correcting for aperiodic activity, we no longer observed a positive relationship between theta activity and WM load as in the original study (Adam et al., 2018). This difference is likely driven by two factors. First, the large, event-related decrease in aperiodic activity is conflated with theta oscillations, which the filter-Hilbert approach cannot disentangle. Second, only 42% of participants had detectable theta oscillations according to spectral parameterization, suggesting very weak, if any, theta oscillations. This is consistent with other research showing theta oscillations are surprisingly scarce within, and across, groups of participants (Bailey et al., 2020; Inanaga, 1998; Mitchell et al., 2008; Voloh & Womelsdorf, 2018; Wilson et al., 2022). We speculate that the ability to detect theta oscillation at the scalp depends on anatomy, such as cortical folding or hippocampal orientation (Hsieh & Ranganath, 2014; Mitchell et al., 2008).

In contrast to theta, almost all participants had detectable alpha oscillations (91%), which might contribute to the finding that alpha oscillations are less susceptible to masking by aperiodic dynamics than theta (Donoghue, Dominguez, et al., 2020). Correcting for aperiodic activity, we found an effect of aperiodic-adjusted alpha power due to WM load and performance. Lateralized effects were only observed during the experiment using a high load only and separating trials based on WM accuracy, in which relative alpha power was more suppressed on the contralateral visual cortex of the attended hemifield. However, the original paper reported much stronger lateralized effects during both the cue and retention period (Adam et al., 2018). Without baselining (subtracting the cue period from the retention period), we do find similarly strong lateralized effects of aperiodic-adjusted alpha power. In summary, alpha oscillations in these datasets were involved in both visual WM and attention mechanisms (D’Esposito & Postle, 2015; Gevins & Smith, 2000; Halgren et al., 2002; Pavlov & Kotchoubey, 2022).

Though our results show aperiodic activity is involved in visual WM, there are methodological limitations to be addressed. First, spectral parameterization on task-based data offers much shorter time windows than resting-state paradigms (Colombo et al., 2019; Helson et al., 2023; McKeon et al., 2024; Smith, Kosik, van Engen, et al., 2023; Waschke et al., 2021), resulting in noisier single-trial power spectra. Although still tractable (Frelih et al., 2024; Höhn et al., 2024; Kałamała et al., 2024; Podvalny et al., 2015; Virtue-Griffiths et al., 2022; Waschke et al., 2021), we minimized noise by rejecting trials based on poor model fits. Another issue became apparent while visually examining power spectra to determine whether participants exhibited oscillatory activity within the theta or alpha band. These canonical frequency bands are adjacent to one another, and some participants exhibited identifiable peaks in both bands, some only in one band, and others in neither. However, there were cases in which a lower center frequency alpha peak bled into the theta range, making it difficult to determine whether a peak should be considered alpha, or theta based on traditional band definitions. This is also a concern for bandpass filters in which neighboring band activity infiltrates the band of interest (Pavlov & Kotchoubey, 2022). Future research could leverage Laplacian filtering or data-driven spatial filters (Nikulin et al., 2011; Schaworonkow & Voytek, 2021) to disentangle these neighboring frequency bands given their high separability in the spatial domain.

Here, we provide correlational evidence that aperiodic activity is dynamic during visual WM. We have also shown that it can mask theta oscillatory dynamics. One way to potentially causally disentangle the role of oscillatory versus aperiodic dynamics in human WM would be to leverage non-invasive neurostimulation. Neurostimulation is commonly used to study the causal role of oscillations, via transcranial Alternating Current Stimulation (tACS) (Grover et al., 2023). Because aperiodic activity consists of arrhythmic fluctuations, transcranial Random Noise Stimulation (tRNS) – white noise stimulation – is a candidate for targeting aperiodic activity, though white noise is different from endogenous aperiodic activity. tRNS flattens spectal power (van Bueren et al., 2023), effectively showing that aperiodic activity can be targeted with non-invasive electrical stimulation. Thus, it may be possible to test the causal role of aperiodic activity in human visual WM using aperiodic stimulation that more closely mimics endogenous aperiodic activity.

While we did not observe load-dependent frontal midline aperiodic changes, we show that previous frontal midline theta results that do not correct for aperiodic activity likely do not reflect theta oscillations. It will be interesting to investigate if our results based on extracranial EEG translate to intracranial EEG and animal models of WM, and if our results are driven by the imprecision of EEG to detect frontal midline theta oscillations. While our results do not invalidate previous research into extracranial theta oscillations in relation to WM, they challenge popular, widely held theories regarding the mechanistic role for theta oscillations to group or segregate channels of information (Lisman & Jensen, 2013). After all, if people reliably perform WM tasks in the absence of detectable theta oscillations, then either our detection ability is severely flawed, or our theories are incomplete (Van Bree et al., 2024).

## Code and Data availability

All data used in this study are publicly available at: https://osf.io/8xuk3/

Adam, K. C. S., Robison, M. K., & Vogel, E. K. (2018, June 25). Open Data 2018: Contralateral delay activity tracks fluctuations in working memory performance. https://doi.org/10.17605/OSF.IO/8XUK3

All code used for all analyses and plots are publicly available on GitHub at: https://github.com/voytekresearch/WM_specparam

## Acknowledgements

We thank Andrew Bender, Dillan Cellier, Ryan Hammonds, Blanca Martin-Burgos, Michael Preston, Eena Kosik, Sydney Smith and Christian Cazares for their advice and feedback on the manuscript. We also want to thank Kirsten Adam, and the Vogel lab for making their data public. Funding was provided by a NIH National Institute of General Medical Sciences grant R01GM134363-01 (to B.V.).

## Author contributions

Q.v.E., K.A., T.D., and B.V. conceived of the experiments and contributed to the intellectual and methodological development of the ideas. Q.v.E., G.C., A.S., and T.D. wrote analysis code and analyzed data. Q.v.E., K.A., T.D., and B.V. wrote and edited the manuscript.

## References

Adam, K. C. S., Robison, M. K., & Vogel, E. K. (2018). Contralateral Delay Activity Tracks Fluctuations in Working Memory Performance. Journal of Cognitive Neuroscience, 30(9), 1229–1240. 10.1162/jocn_a_01233

Adam, K. C. S., Vogel, E. K., & Awh, E. (2020). Multivariate analysis reveals a generalizable human electrophysiological signature of working memory load. Psychophysiology, 57(12), e13691. 10.1111/psyp.13691

Akbarian, F., Rossi, C., Costers, L., D’hooghe, M. B., D’haeseleer, M., Nagels, G., & Schependom, J. V. (2023). Stimulus-related modulation in the 1/f spectral slope suggests an impaired inhibition of irrelevant information in people with multiple sclerosis (p. 2023.12.28.573572). bioRxiv. 10.1101/2023.12.28.573572

Baddeley, A. (1992). Working Memory. Science, 255(5044), 556.

Bae, G.-Y., & Luck, S. J. (2018). Dissociable Decoding of Spatial Attention and Working Memory from EEG Oscillations and Sustained Potentials. Journal of Neuroscience, 38(2), 409–422. 10.1523/JNEUROSCI.2860-17.2017

Bailey, N. W., Freedman, G., Raj, K., Spierings, K. N., Piccoli, L. R., Sullivan, C. M., Chung, S. W., Hill, A. T., Rogasch, N. C., & Fitzgerald, P. B. (2020). Mindfulness Meditators Show Enhanced Accuracy and Different Neural Activity During Working Memory. Mindfulness, 11(7), 1762–1781. 10.1007/s12671-020-01393-8

Bender, A., Zhao, C., Vogel, E., Awh, E., & Voytek, B. (2025). Differential representations of spatial location by aperiodic and alpha oscillatory activity in working memory (p. 2025.03.21.644412). bioRxiv. 10.1101/2025.03.21.644412

Berger, B., & Sauseng, P. (2022). Brain rhythms: How control gets into working memory. Current Biology, 32(10), R479–R481. 10.1016/j.cub.2022.04.036

Brake, N., Duc, F., Rokos, A., Arseneau, F., Shahiri, S., Khadra, A., & Plourde, G. (2024). A neurophysiological basis for aperiodic EEG and the background spectral trend. Nature Communications, 15(1), 1514. 10.1038/s41467-024-45922-8

Cole, S., Donoghue, T., Gao, R., & Voytek, B. (2019). NeuroDSP: A package for neural digital signal processing. Journal of Open Source Software, 4(36), 1272. 10.21105/joss.01272

Colombo, M. A., Napolitani, M., Boly, M., Gosseries, O., Casarotto, S., Rosanova, M., Brichant, J.-F., Boveroux, P., Rex, S., Laureys, S., Massimini, M., Chieregato, A., & Sarasso, S. (2019). The spectral exponent of the resting EEG indexes the presence of consciousness during unresponsiveness induced by propofol, xenon, and ketamine. NeuroImage, 189, 631–644. 10.1016/j.neuroimage.2019.01.024

Compte, A., Brunel, N., Goldman-Rakic, P. S., & Wang, X.-J. (2000). Synaptic Mechanisms and Network Dynamics Underlying Spatial Working Memory in a Cortical Network Model. Cerebral Cortex, 10(9), 910–923. 10.1093/cercor/10.9.910

Cowan, N. (2010). The Magical Mystery Four: How is Working Memory Capacity Limited, and Why? Current Directions in Psychological Science, 19(1), 51–57. 10.1177/0963721409359277

Cunningham, E., Zimnicki, C., & Beck, D. M. (2023). The Influence of Prestimulus 1/f-Like versus Alpha-Band Activity on Subjective Awareness of Auditory and Visual Stimuli. The Journal of Neuroscience, 43(37), 6447–6459. 10.1523/JNEUROSCI.0238-23.2023

D’Esposito, M., & Postle, B. R. (2015). The Cognitive Neuroscience of Working Memory. Annual Review of Psychology, 66(1), 115–142. 10.1146/annurev-psych-010814-015031

Donoghue, T. (2024). A systematic review of aperiodic neural activity in clinical investigations. 10.1101/2024.10.14.24314925

Donoghue, T., Dominguez, J., & Voytek, B. (2020). Electrophysiological Frequency Band Ratio Measures Conflate Periodic and Aperiodic Neural Activity. eNeuro, 7(6). 10.1523/ENEURO.0192-20.2020

Donoghue, T., Haller, M., Peterson, E. J., Varma, P., Sebastian, P., Gao, R., Noto, T., Lara, A. H., Wallis, J. D., Knight, R. T., Shestyuk, A., & Voytek, B. (2020). Parameterizing neural power spectra into periodic and aperiodic components. Nature Neuroscience, 23(12), 1655–1665. 10.1038/s41593-020-00744-x

Foster, J. J., Sutterer, D. W., Serences, J. T., Vogel, E. K., & Awh, E. (2016). The topography of alpha-band activity tracks the content of spatial working memory. Journal of Neurophysiology, 115(1), 168–177. 10.1152/jn.00860.2015

Frelih, T., Matkovič, A., Mlinarič, T., Bon, J., & Repovš, G. (2024). Modulation of aperiodic EEG activity provides sensitive index of cognitive state changes during working memory task. eLife, 13. 10.7554/eLife.101071.1

Fukuda, K., Kang, M.-S., & Woodman, G. F. (2016). Distinct neural mechanisms for spatially lateralized and spatially global visual working memory representations. Journal of Neurophysiology, 116(4), 1715–1727. 10.1152/jn.00991.2015

Fukuda, K., Mance, I., & Vogel, E. K. (2015). □ Power Modulation and Event-Related Slow Wave Provide. The Journal of Neuroscience, 35(41), 14009–14016.

Gao, R., Peterson, E. J., & Voytek, B. (2017). Inferring synaptic excitation/inhibition balance from field potentials. NeuroImage, 158, 70–78. 10.1016/j.neuroimage.2017.06.078

Gevins, A., & Smith, M. E. (2000). Neurophysiological Measures of Working Memory and Individual Differences in Cognitive Ability and Cognitive Style. Cerebral Cortex, 10(9), 829–839. 10.1093/cercor/10.9.829

Grover, S., Fayzullina, R., Bullard, B. M., Levina, V., & Reinhart, R. M. G. (2023). A meta-analysis suggests that tACS improves cognition in healthy, aging, and psychiatric populations. Science Translational Medicine, 15(697), eabo2044. 10.1126/scitranslmed.abo2044

Gyurkovics, M., Clements, G. M., Low, K. A., Fabiani, M., & Gratton, G. (2022). Stimulus-Induced Changes in 1/f-like Background Activity in EEG. Journal of Neuroscience, 42(37), 7144–7151. 10.1523/JNEUROSCI.0414-22.2022

Halgren, E., Boujon, C., Clarke, J., Wang, C., & Chauvel, P. (2002). Rapid Distributed Fronto-parieto-occipital Processing Stages During Working Memory in Humans. Cerebral Cortex, 12(7), 710–728. 10.1093/cercor/12.7.710

He, B. J. (2014). Scale-free brain activity: Past, present, and future. Trends in Cognitive Sciences, 18(9), 480–487. 10.1016/j.tics.2014.04.003

Helson, P., Lundqvist, D., Svenningsson, P., Vinding, M. C., & Kumar, A. (2023). Cortex-wide topography of 1/f-exponent in Parkinson’s disease. Npj Parkinson’s Disease, 9(1), Article 1. 10.1038/s41531-023-00553-6

Herweg, N. A., Solomon, E. A., & Kahana, M. J. (2020). Theta Oscillations in Human Memory. Trends in Cognitive Sciences, 24(3), 208–227. 10.1016/j.tics.2019.12.006

Höhn, C., Hahn, M. A., Lendner, J. D., & Hoedlmoser, K. (2024). Spectral Slope and Lempel–Ziv Complexity as Robust Markers of Brain States during Sleep and Wakefulness. eNeuro, 11(3). 10.1523/ENEURO.0259-23.2024

Hsieh, L.-T., & Ranganath, C. (2014). Frontal midline theta oscillations during working memory maintenance and episodic encoding and retrieval. NeuroImage, 85, 721–729. 10.1016/j.neuroimage.2013.08.003

Hunter, J. D. (2007). Matplotlib: A 2D graphics environment. Computing in Science & Engineering, 9(3), 90–95. 10.1109/MCSE.2007.55

Inanaga, K. (1998). Frontal midline theta rhythm and mental activity. Psychiatry and Clinical Neurosciences, 52(6), 555–566. 10.1046/j.1440-1819.1998.00452.x

Jensen, O., & Tesche, C. D. (2002). Frontal theta activity in humans increases with memory load in a working memory task: Frontal theta increases with memory load. European Journal of Neuroscience, 15(8), 1395–1399. 10.1046/j.1460-9568.2002.01975.x

Jones, S. R. (2016). When brain rhythms aren’t ‘rhythmic’: Implication for their mechanisms and meaning. Current Opinion in Neurobiology, 40, 72–80. 10.1016/j.conb.2016.06.010

Kałamała, P., Gyurkovics, M., Bowie, D. C., Clements, G. M., Low, K. A., Dolcos, F., Fabiani, M., & Gratton, G. (2024). Event-induced modulation of aperiodic background EEG: Attention-dependent and age-related shifts in E:I balance, and their consequences for behavior. Imaging Neuroscience, 2, 1–18. 10.1162/imag_a_00054

Karalunas, S. L., Ostlund, B. D., Alperin, B. R., Figuracion, M., Gustafsson, H. C., Deming, E. M., Foti, D., Antovich, D., Dude, J., Nigg, J., & Sullivan, E. (2022). Electroencephalogram aperiodic power spectral slope can be reliably measured and predicts ADHD risk in early development. Developmental Psychobiology, 64(3), e22228. 10.1002/dev.22228

Klimesch, W. (1999). EEG alpha and theta oscillations reflect cognitive and memory performance: A review and analysis. Brain Research Reviews, 29(2–3), 169–195. 10.1016/S0165-0173(98)00056-3

Lara, A. H., & Wallis, J. D. (2015). The Role of Prefrontal Cortex in Working Memory: A Mini Review. Frontiers in Systems Neuroscience, 9, 173. 10.3389/fnsys.2015.00173

Lendner, J. D., Harler, U., Daume, J., Engel, A. K., Zöllner, C., Schneider, T. R., & Fischer, M. (2023). Oscillatory and aperiodic neuronal activity in working memory following anesthesia. Clinical Neurophysiology, 150, 79–88. 10.1016/j.clinph.2023.03.005

Lendner, J. D., Helfrich, R. F., Mander, B. A., Romundstad, L., Lin, J. J., Walker, M. P., Larsson, P. G., & Knight, R. T. (2020). An electrophysiological marker of arousal level in humans. eLife, 9, e55092. 10.7554/eLife.55092

Lisman, J. E., & Jensen, O. (2013). The Theta-Gamma Neural Code. Neuron, 77(6), 1002–1016. 10.1016/j.neuron.2013.03.007

Lu, L., Gao, Z., Wei, Z., & Yi, M. (2023). Working memory depends on the excitatory–inhibitory balance in neuron–astrocyte network. Chaos: An Interdisciplinary Journal of Nonlinear Science, 33(1), 013127. 10.1063/5.0126890

McKeon, S. D., Perica, M. I., Parr, A. C., Calabro, F. J., Foran, W., Hetherington, H., Moon, C.-H., & Luna, B. (2024). Aperiodic EEG and 7T MRSI evidence for maturation of E/I balance supporting the development of working memory through adolescence. Developmental Cognitive Neuroscience, 66, 101373. 10.1016/j.dcn.2024.101373

Medendorp, W. P., Kramer, G. F. I., Jensen, O., Oostenveld, R., Schoffelen, J.-M., & Fries, P. (2007). Oscillatory activity in human parietal and occipital cortex shows hemispheric lateralization and memory effects in a delayed double-step saccade task. Cerebral Cortex (New York, N.Y.: 1991), 17(10), 2364–2374. 10.1093/cercor/bhl145

Miller, G. A. (1956). The Magical Number Seven, Plus or Minus Two Some Limits on Our Capacity for Processing Information. Psychological Review, 63(2), 81.

Mitchell, D. J., McNaughton, N., Flanagan, D., & Kirk, I. J. (2008). Frontal-midline theta from the perspective of hippocampal “theta.” Progress in Neurobiology, 86(3), 156–185. 10.1016/j.pneurobio.2008.09.005

Nikulin, V. V., Nolte, G., & Curio, G. (2011). A novel method for reliable and fast extraction of neuronal EEG/MEG oscillations on the basis of spatio-spectral decomposition. NeuroImage, 55(4), 1528–1535. 10.1016/j.neuroimage.2011.01.057

Oberauer, K., Farrell, S., Jarrold, C., & Lewandowsky, S. (2016). What limits working memory capacity? Psychological Bulletin, 142(7), 758–799. 10.1037/bul0000046

Palva, S., & Palva, J. M. (2007). New vistas for α-frequency band oscillations. Trends in Neurosciences, 30(4), 150–158. 10.1016/j.tins.2007.02.001

Pavlov, Y. G., & Kotchoubey, B. (2022). Oscillatory brain activity and maintenance of verbal and visual working memory: A systematic review. Psychophysiology, 59(5), e13735. 10.1111/psyp.13735

Podvalny, E., Noy, N., Harel, M., Bickel, S., Chechik, G., Schroeder, C. E., Mehta, A. D., Tsodyks, M., & Malach, R. (2015). A unifying principle underlying the extracellular field potential spectral responses in the human cortex. Journal of Neurophysiology, 114(1), 505–519. 10.1152/jn.00943.2014

Sauseng, P., Klimesch, W., Heise, K. F., Gruber, W. R., Holz, E., Karim, A. A., Glennon, M., Gerloff, C., Birbaumer, N., & Hummel, F. C. (2009). Brain Oscillatory Substrates of Visual Short-Term Memory Capacity. Current Biology, 19(21), 1846–1852. 10.1016/j.cub.2009.08.062

Schaworonkow, N., & Voytek, B. (2021). Enhancing oscillations in intracranial electrophysiological recordings with data-driven spatial filters. PLOS Computational Biology, 17(8), e1009298. 10.1371/journal.pcbi.1009298

Smith, S. E., Kosik, E. L., van Engen, Q., Kohn, J., Hill, A. T., Zomorrodi, R., Blumberger, D. M., Daskalakis, Z. J., Hadas, I., & Voytek, B. (2023). Magnetic seizure therapy and electroconvulsive therapy increase aperiodic activity. Translational Psychiatry, 13(1), Article 1. 10.1038/s41398-023-02631-y

Smith, S. E., Ma, V., Gonzalez, C., Chapman, A., Printz, D., Voytek, B., & Soltani, M. (2023). Clinical EEG slowing induced by electroconvulsive therapy is better described by increased frontal aperiodic activity. Translational Psychiatry, 13(1), Article 1. 10.1038/s41398-023-02634-9

Soltani Zangbar, H., Ghadiri, T., Seyedi Vafaee, M., Ebrahimi Kalan, A., Fallahi, S., Ghorbani, M., & Shahabi, P. (2020). Theta Oscillations Through Hippocampal/Prefrontal Pathway: Importance in Cognitive Performances. Brain Connectivity, 10(4), 157–169. 10.1089/brain.2019.0733

Vallat, R. (2018). Pingouin: Statistics in Python. Journal of Open Source Software, 3(31), 1026. 10.21105/joss.01026

Van Bree, S., Levenstein, D., Krause, M., Voytek, B., & Gao, R. (2024). Decoupling Measurements and Processes: On the Epiphenomenon Debate Surrounding Brain Oscillations in Field Potentials. 10.31234/osf.io/knjfw

van Bueren, N. E. R. van, Ven, S. H. G. van der, Hochman, S., Sella, F., & Kadosh, R. C. (2023). Human neuronal excitation/inhibition balance explains and predicts neurostimulation induced learning benefits. PLOS Biology, 21(8), e3002193. 10.1371/journal.pbio.3002193

van der Werf, J., Jensen, O., Fries, P., & Medendorp, W. P. (2008). Gamma-Band Activity in Human Posterior Parietal Cortex Encodes the Motor Goal during Delayed Prosaccades and Antisaccades. Journal of Neuroscience, 28(34), 8397–8405. 10.1523/JNEUROSCI.0630-08.2008

van Dijk, H., van der Werf, J., Mazaheri, A., Medendorp, W. P., & Jensen, O. (2010). Modulations in oscillatory activity with amplitude asymmetry can produce cognitively relevant event-related responses. Proceedings of the National Academy of Sciences, 107(2), 900–905. 10.1073/pnas.0908821107

Vanderwolf, C. H. (1969). Hippocampal electrical activity and voluntary movement in the rat. Electroencephalography and Clinical Neurophysiology, 26(4), 407–418. 10.1016/0013-4694(69)90092-3

Virtue-Griffiths, S., Fornito, A., Thompson, S., Biabani, M., Tiego, J., Thapa, T., & Rogasch, N. C. (2022). Task-related changes in aperiodic activity are related to visual working memory capacity independent of event-related potentials and alpha oscillations [Preprint]. Neuroscience. 10.1101/2022.01.18.476852

Vogel, E. K., & Machizawa, M. G. (2004). Neural activity predicts individual differences in visual working memory capacity. Nature, 428(6984), 748–751. 10.1038/nature02447

Voloh, B., & Womelsdorf, T. (2018). Cell-Type Specific Burst Firing Interacts with Theta and Beta Activity in Prefrontal Cortex During Attention States. Cerebral Cortex, 28(12), 4348–4364. 10.1093/cercor/bhx287

Voytek, B., Kramer, M. A., Case, J., Lepage, K. Q., Tempesta, Z. R., Knight, R. T., & Gazzaley, A. (2015). Age-Related Changes in 1/f Neural Electrophysiological Noise. Journal of Neuroscience, 35(38), 13257–13265. 10.1523/JNEUROSCI.2332-14.2015

Wang, X.-J. (1999). Synaptic Basis of Cortical Persistent Activity: The Importance of NMDA Receptors to Working Memory. Journal of Neuroscience, 19(21), 9587–9603. 10.1523/JNEUROSCI.19-21-09587.1999

Waschke, L., Donoghue, T., Fiedler, L., Smith, S., Garrett, D. D., Voytek, B., & Obleser, J. (2021). Modality-specific tracking of attention and sensory statistics in the human electrophysiological spectral exponent. eLife, 10, e70068. 10.7554/eLife.70068

Waskom, M. L. (2021). Journal of Open Source Software, 6(60), 3021. 10.21105/joss.03021

Wilson, L. E., da Silva Castanheira, J., & Baillet, S. (2022). Time-resolved parameterization of aperiodic and periodic brain activity. eLife, 11, e77348. 10.7554/eLife.77348

